# Effects of age and dietary methionine restriction on cognitive and behavioural phenotypes in the rTg4510 model of frontotemporal dementia

**DOI:** 10.1101/2024.03.05.583295

**Authors:** Marina Souza Matos, Annesha Sil, Gernot Riedel, Bettina Platt, Mirela Delibegovic

## Abstract

Metabolic disorders such as diabetes and obesity are linked to neurodegenerative diseases, with evidence of decreased brain glucose metabolism and insulin resistance in patients with dementia. Given the rising prevalence of age-related diseases, lifestyle adjustments and nutritional interventions are gaining interest. Dietary methionine restriction (MR) is a nutritional intervention that enhances insulin sensitivity and delays ageing-associated metabolic alterations. Since the potential impact of MR on neurodegenerative diseases like dementia is not fully understood, we here examined the metabolic and behavioural phenotypes of a murine tauopathy model (rTg4510), which overexpresses human P301L mutated tau, and assessed the impact of an 8-week dietary MR. The rTg4510 mice and wild type (WT) littermates were assessed at 6 and 12 months of age. While rTg4510 mice displayed progressive behavioural and motor impairments at 6 and 12 months of age, MR led to significant benefits in the aged 12-month-old cohort, improving motor coordination and learning, short-term memory, and social recognition. These effects were accompanied by increased glycolysis in the hippocampus and higher FGF21 levels in the cortex. These benefits occurred in the absence of alterations in glucose metabolism/adiposity in this model. Overall, our results support the positive impact of MR on rTg4510 mice, suggesting this as a potential therapeutic intervention to delay and/or improve the progression in tau-related disease.

## Introduction

Neurodegenerative diseases are a major societal concern caused by genetic, environmental, and lifestyle-related factors. Ageing is the main risk factor for associated disorders such as dementia (1), a clinical syndrome with progressive cognitive decline (2). Its presentation varies depending on the subtype (3), e.g. Alzheimer’s Disease (AD), vascular dementia, Lewy body dementia and frontotemporal dementia (FTD) (4). As one of the most prevalent tauopathies characterised by abnormal accumulation of tau protein in the brain, FTD accounts for 10 – 20% of all cases (5). Clinical signs categorise symptoms, e.g. behavioural variant type of FTD (bvFTD) exhibits personality changes, disinhibition, altered dietary preferences, and appetite shifts (6,7).

Although FTD symptoms are well described, the mechanisms behind its pathophysiology remain to be elucidated. Mutations of the tau gene, which encodes the microtubule-associated protein tau (MAPT), have been linked with familial forms (8). These promote abnormal hyperphosphorylation of tau, aggregation of misfolded protein and extracellular neurofibrillary tangles (NFT) deposition (9). However, most FTD cases are sporadic and associated with modifiable risk factors, including the prevalence of cardiovascular diseases (CVD) and type 2 diabetes (T2D), amongst others (10).

Increased prevalence of age-related diseases increases interest in finding nutritional interventions that could delay the onset or progression of these disorders (11). Methionine restriction (MR) has shown benefits in metabolism and longevity, by mimicking caloric restriction (CR) without promoting malnutrition. MR reduced cognitive decline and improved glucose homeostasis in mice (12–14), aided the reduction in adiposity and obesity, and reversed age-related metabolic disruption, resulting in a ‘younger’ metabolic phenotype (15). However, it is unclear whether the beneficial effects of MR extend to dementia and other neurodegenerative diseases.

An essential contributor to the benefits of MR in longevity and metabolic health is fibroblast growth factor 21 (FGF21), predominantly synthesised by the liver but also expressed in the brain (16–19). FGF21 treatment has shown neuroprotective effects and improved cognition in obese, insulin-resistant rats (20). Despite conflicting findings on FGF21 gene expression in the hippocampus, it is crucial for MR-related cognitive improvements (21,22). Moreover, MR in aged mice also promoted memory enhancement alongside increased FGF21 levels (23).

Given the benefits of MR on metabolic and cognitive health and disturbances in brain glucose homeostasis seen in dementia patients (9,24,25), this study investigated behavioural, molecular, and metabolic effects in a mouse model of FTD. We carried out a longitudinal MR study in the well-characterised rTg4510 tauopathy model at 6– and 12 months of age. rTg4510 mice develop various FTD-like features as they age driven by severe tau pathology (26) and exacerbated in female vs male mice (27,28). We therefore assessed the efficacy of 8 weeks of MR in female mice.

## Results

### MR diet did not alter body composition in rTg4510 mice

Cognitive decline and metabolic disturbances are highly associated with ageing. Thus, we examined the effects of tau, age, and diet on body composition. Total body weight of WT and rTg4510 mice did not differ between 6– and 12-months groups, but an overall genotype effect was revealed when comparing groups at 12 months (F (1, 26) = 5.318, p<0.05, Fig. 1A), i.e., rTg4510s weighed less than their littermate controls regardless of diet (Fig. 1A). 12-month-old WT mice had more fat and lean mass than age-matched rTg4510 animals (Fig. 1B & C), yet WT mice on MR diet displayed lower fat content compared to control diet. No dietary effect was seen in fat or lean mass in rTg4510 mice (Fig. 1B & C).

**FIGURE 1:**
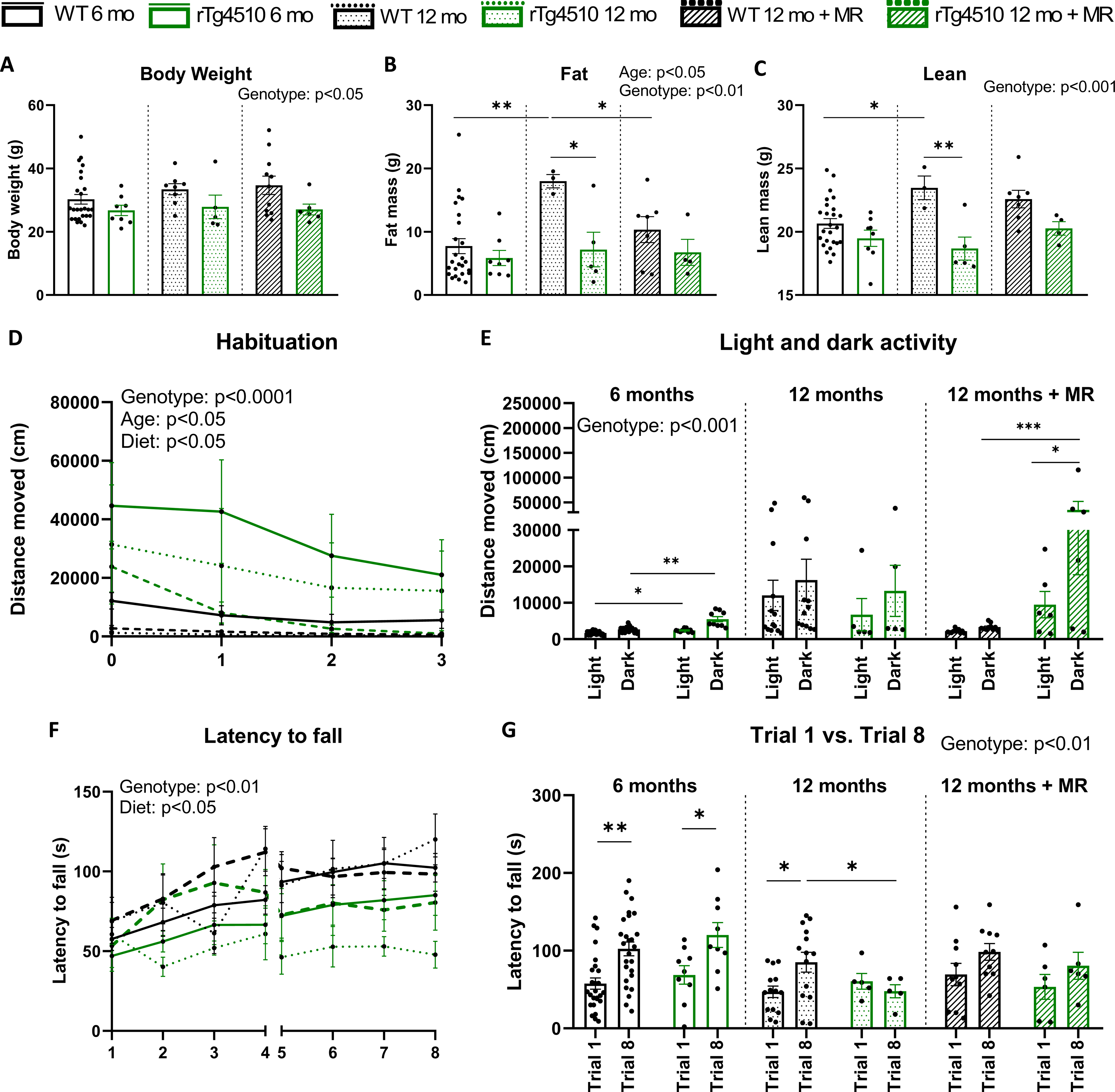
Motor activity, performance, learning, and body composition. (A) Body weight, (B) fat, and (C) lean mass determined by Echo MRI scans (n = 25 (WT 6mo), n = 8 (rTg 6mo), n = 3 or 8 (WT 12mo), n = 7 or 11 (WT 12mo MR), n = 5 (rTg 12mo), n = 4 (rTg 12mo MR)). (D) Distance moved (cm/hour) during 3h-habituation period. (E) Average distance moved during light and dark periods. (F) Latency to fall (seconds) from the RotaRod. (G) Motor learning determined by the latency to fall (seconds) from the rod over trial 1 and trial 8 (n = 26 (WT 6mo), n = 14 (rTg 6mo), n = 9 (WT 12mo), n = 10 (WT 12mo MR), n = 5 (rTg 12mo), n = 6 (rTg 12mo MR)). Age (WT vs rTg 6 months) and diet (control vs MR 12 months) analyses were performed separately by 3-way ANOVA followed by Tukey’s multiple comparison test, or 2-way ANOVA followed by Bonferroni’s multiple comparison test. Data were plotted together for visualisation purposes. Habituation data analysed by nonlinear regression (one-phase decay), followed by 3-way ANOVA. Data are presented as scatter plus mean ± SEM. *p<0.05, **p<0.01, ***p<0.001, ****p<0.0001.

Food and water seeking behaviour over one week of PhenoTyper testing was measured (Fig. S2). Overall, both WT and rTg4510 mice spent more time in the food zone at 12 months of age compared to 6 months. A genotype effect in both times spent in water zone indicate that rTg4510 mice spent more time in both zones at both ages. Dietary group analysis also confirmed previous activity observations, i.e., increased rTg4510 mice activity during light and dark phases in both zones (Supplementary Table 1 for full statistics). No dietary effect was observed as such. Overall, despite rTg4510 mice spending longer in food and water zone than WT mice, this behaviour is not reflected in changes on body weight. Interestingly, WT animals on MR diet still present less fat content compared to their 12-month-control littermates, despite no changes in eating behaviour.

### MR diet improves habituation to a new environment and motor coordination in 12-month-old rTg4510 mice

Hyperactivity is a typical trait of the rTg4510 model, mimicking FTD patients’ symptoms (29). In line with previous data, we detected a significant difference in motor activity between the genotypes during habituation (3 hrs) to a novel environment in PhenoTyper cages. rTg4510 mice were more active than WT at both 6– and 12-months during habituation (F (1, 187) = 27.01, p<0.0001, age vs. genotype analysis, Fig. 1D). An overall diet effect (F (1, 111) = 4.758, p<0.05) and a genotype x diet interaction (F (1, 111) = 6.157, p<0.05) occurred, revealing that MR had different effects on the animals’ activity depending on genotype during habituation. Throughout the habituation period, MR decreased the hyperactivity of rTg4510 mice compared to rTg4510 on control diet, reaching similar activity levels to WT mice after 3 hours. In comparison, rTg4510 mice on control diet remained hyperactive during the habituation period (Fig 1D). In control mice, MR increased activity during habituation but reached a similar ambulatory level as the control diet group. Overall, this suggests that MR decreased rTg4510 mice activity during habituation to an unfamiliar environment. Separate WT habituation graphs are shown in S3 for visualisation purposes.

After habituation, 6-month-old rTg4510 mice remained significantly more active than WTs during both, dark and light phases on control diet (Fig. 1E), suggesting that habituation had not normalised the innate hyperactive phenotype of rTg4510 mice. Age analysis did not reveal statistical differences between 6– and 12-months old mice; however, rTg4510 animals remained hyperactive at older age (genotype effect: F (1, 47) = 9.007, p<0.01). The average hourly activity during both day and night periods did not indicate a dietary effect, but it is worth noting that the MR effect on rTg4510 behavioural activity was different in the novel vs familiar environment. Overall, these data suggest intact basic circadian and ultradian patterns, but higher locomotor activity of rTg4510 mice vs WT animals.

Next, motor coordination and learning were assessed over 2 consecutive days using the RotaRod. rTg4510 mice spent less time on the rod compared to the WT group at 6– and 12-months of age (latency to fall over all trials, Fig. 1F), indicative of a motor strength and/or coordination impairment. However, rTg4510 mice on MR were able to stay on longer compared to rTg4510 on the control diet (diet effect: F (1, 208) = 6.309, p<0.05). No such diet effect was detected in WT mice. This was confirmed further by a genotype x diet interaction (F (1, 208) = 4.186, p<0.05), suggesting that MR improves motor coordination in transgenic mice only. Moreover, a significant improvement over trials was revealed (time on the rod) for both groups at 6 and 12 months of age on control diet (trial 1 vs 8, F (1, 50) = 13.62, p<0.001, Fig. 1G), hence, motor learning was intact in both genotypes. Still, at 12 months of age, a reliable genotype effect (F (1, 50) = 7.498, p<0.01, Fig. 1I) was observed, indicative of a motor learning deficit in older rTg4510 on control diet. Overall, these data indicate that rTg4510 mice exhibit motor impairments at 12 months of age, and MR improves motor coordination specifically in transgenic mice.

### MR rescues short-term spatial memory, improves between-trial habituation, and social recognition in 12-month-old rTg4510 mice

To investigate spatial (short-term) working memory, we determined the alternation index (in %) during the Y-maze task (Fig. 2A). A typical correct alternation score of 50–60% has been previously observed in WT mice at 6 months (30). Here, correct and incorrect alternations did not differ between WT and rTg4510 mice (Fig. 2B). Within the same genotype, both WT and rTg4510 mice performed more correct than incorrect alternations, indicating intact spatial working memory at this age. However, at 12 months, both WT and rTg4510 mice displayed correct and incorrect alternations equally, indicating a worsening of short-term spatial memory with age. Remarkably, this deficit was rescued by MR, as both MR groups performed more correct than incorrect alternations (Fig. 2B). The greatest difference was seen in MR rTg4510 mice (p<0.0001; Fig. 2B), indicating that the benefits of MR were more prominent in the FTD model.

**FIGURE 2:**
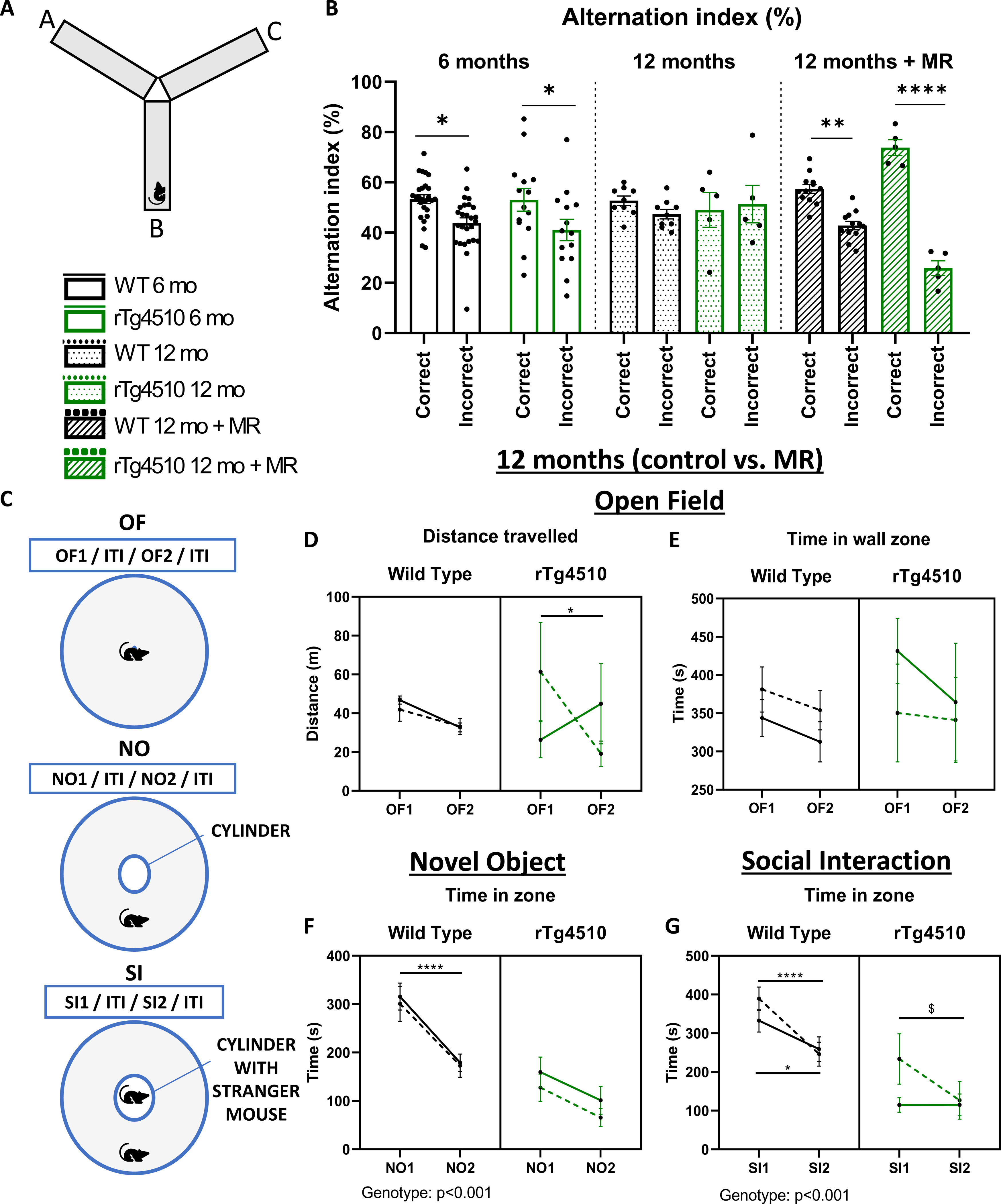
Y-maze and Open Field-Novel Object-Social Interaction paradigm. (A) Y maze apparatus representation. (B) Alternation index between correct and incorrect alternations (n = 26 (WT 6mo), n = 14 (rTg 6mo), n = 9 (WT 12mo), n = 12 (WT 12mo MR), n = 5 (rTg 12mo), n = 5 (rTg 12mo MR)). Descriptive analyses are relative to diet analyses. No statistical age effect was observed in the Y maze. Age (WT vs rTg 6 months) and diet (control vs MR 12 months) analyses were performed separately by 3-way ANOVA followed by Tukey’s multiple comparison test. Data were plotted together for visualisation purposes and presented as scatter plus mean ± SEM. *p<0.05, **p<0.01, ***p<0.001, ****p<0.0001. (C) Schematic representation of the OF-NO-SI experimental design. (D) Total distance travelled and (E) time spent in the wall zone during trials 1 (OF1) and 2 (OF2) of OF. *p<0.05 rTg4510 MR OF1 vs. OF2. (F) Time spent in the cylinder zone during trials 1 and 2 of NO. ****p<0.0001 WT control and MR NO1 vs. NO2. (G) Time spent in the cylinder/stranger mouse zone during trials 1 and 2 of SI. *p<0.05 WT control SI1 vs. SI2, ****p<0.0001 WT MR SI1 vs. SI2, $p<0.05 rTg4510 MR SI1 vs. SI2. (n = 9 (WT 12mo), n = 12 (WT 12mo MR), n = 5 (rTg 12mo), n = 6 (rTg 12mo MR)). Analyses were performed by 3-way ANOVA followed by Tukey’s multiple comparison test or by Mixed-effects analysis followed by Bonferroni’s multiple comparison test (D and G). Data are presented as scatter plus mean ± SEM.

Next, the behaviour of WT and rTg4510 animals in a novel (trial 1) and a familiar (trial 2) environment was evaluated using the Open Field-Novel Object-Social Interaction (OF-NO-SI) task (Fig. 2C). This task was only performed at 12 months of age. During OF trials, total distance moved was analysed as a measure for exploratory behaviour. Analysis revealed a trial x diet interaction (F (1, 27) = 4.653, p<0.05) and a trial x genotype x diet interaction (F (1, 27) = 6.743, p<0.05, Fig. 2D). This suggests that MR significantly decreased the motor activity of rTg4510 during OF2 compared to OF1 (p<0.05), indicative of improved habituation to the arena (Fig. 2D). A decline in exploration was not seen in rTg4510 mice on control diet, and WT mice on either diet only showed a minor, non-significant decline (Fig. 2D). Time spent in the wall zone was also investigated to examine thigmotaxis and anxiety-like behaviour (31,32). This parameter did not differ between dietary groups (Fig. 2E).

Object exploration was assessed during the novel object (NO, cylinder) segment of the task. Time spent in the object zone during trials 1 and 2 was analysed. A noticeable overall genotype effect was observed for both trials (F (1, 28) = 20.40, p<0.001), as rTg4510 animals spent less time in the NO interaction zone (Fig. 2F). A trial effect (F (1, 28) = 49.86, p<0.0001) and trial x genotype interaction (F (1, 28) = 7.063, p<0.05) were also detected, due to the significant reduction in the time that WT mice spent in the zone during NO2 vs NO1 in both dietary groups (Fig. 2F). The diet did not affect this parameter in either group.

For social interaction (SI) and engagement with a conspecific, an age-and sex-matched stranger mouse was placed inside the cylinder. The between-trials comparison revealed that WT mice on both diets spent less time in the interaction zone during SI2 vs. SI1 (p<0.05 WT control SI1 vs SI2; p<0.0001 WT MR SI1 vs SI2), suggesting recognition of the stranger mouse during the second trial (Fig. 2G). For rTg4510, only mice on MR exhibited a decrease in the time spent in the SI zone between trials, indicating an MR effect on social recognition depending on genotype (p<0.05 rTg MR SI1 vs SI2) (Fig. 2G). Overall trial (F (1, 27) = 26.72, p<0.0001) and genotype effects (F (1, 27) = 20.64, p<0.001) confirmed these observations, together with a trial x diet interaction (F (1, 27) = 7.951, p<0.01, Fig. 2G).

### MR alters tau levels in the brains of 12-month-old rTg4510 mice

Levels of human Tau (hTau) and endogenous mouse Tau (mTau) were determined in the hippocampus of the rTg4510 mice by qPCR. In line with previous data (26,29), a ∼20-fold higher expression of hTau was found in the rTg4510 at both ages, confirming hTau expression in rTg4510 mice (genotype effect F (1, 27) = 138.8, p<0.0001; Fig. 3A). Surprisingly, there were significantly higher hTau mRNA levels in MR rTg4510 mice compared to rTg4510 on control diet (diet effect F (1, 27) = 6.911, p<0.05; Fig. 3A). mTau gene levels were confirmed to be unaffected and equivalent in all studied groups (Fig. 3B).

**FIGURE 3:**
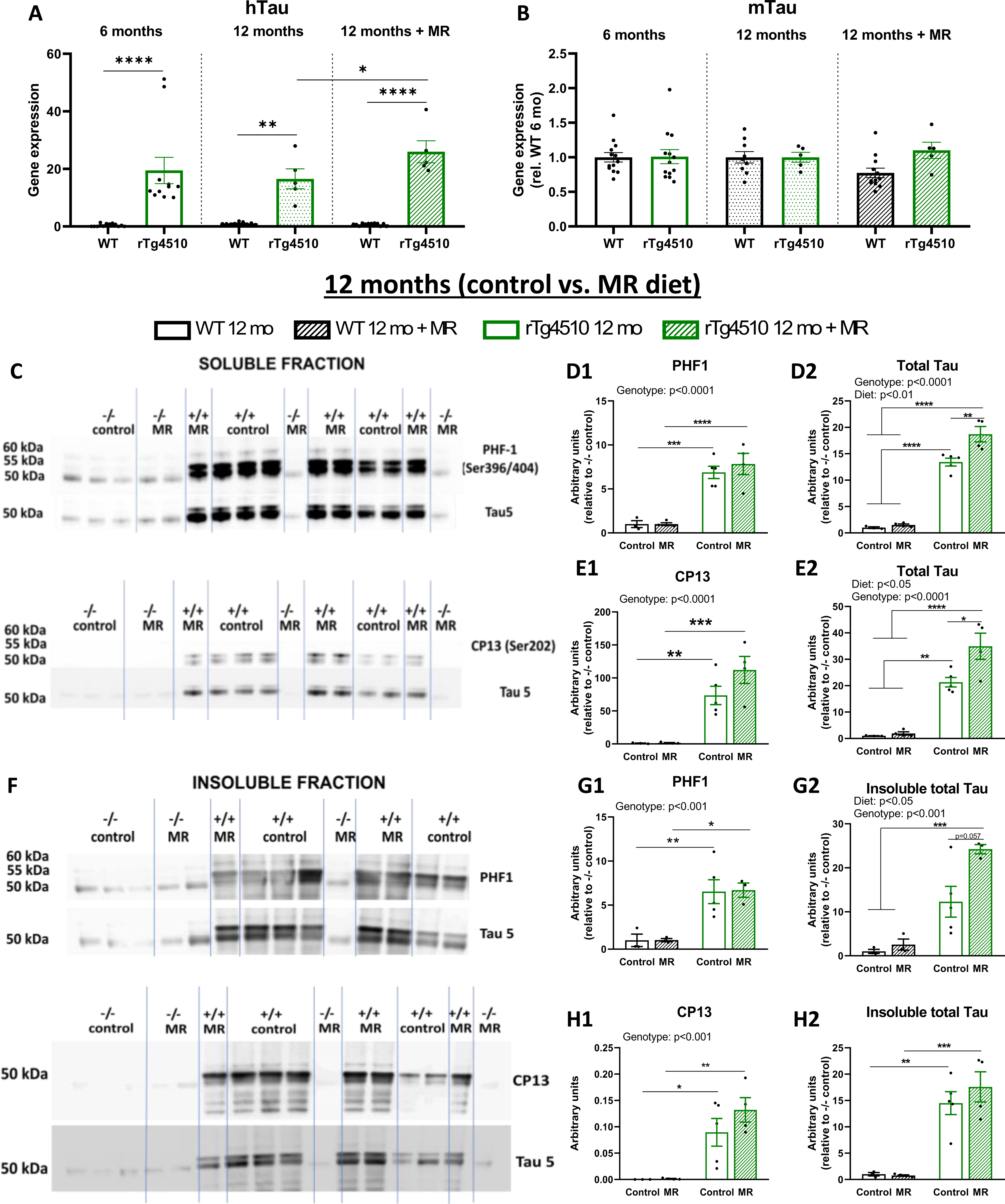
Tau expression. (A) Gene expression of human Tau and (B) mouse Tau in the hippocampus YWHAZ was used as housekeeping gene. (n = 12 (WT 6mo), n = 11-12 (rTg 6mo), n = 9 (WT 12mo), n = 12 (WT 12mo MR), n = 5 (rTg 12mo), n = 5 (rTg 12mo MR)). Data analysed by 2-way ANOVA followed by Bonferroni’s multiple comparison test. (C) Representative immunoblots of soluble phosphorylated PHF1 (Ser396/404), CP13 (Ser 202), and Tau 5 (total Tau) of WT and rTg4510 mice on control or MR diet. (D1) Densitometry analysis of total soluble PHF1, (E1) CP13, and (D2, E2) total Tau. (F) Representative immunoblots of insoluble phosphorylated PHF1 (Ser396/404), CP13 (Ser 202), and Tau 5 (total Tau). (G1) Densitometry analysis of total insoluble PHF1, (H1) CP13, and (G2, H2) total Tau. All data are normalised by their respective Ponceau staining and shown as relative to the group WT (-/-) control diet. Data analysed by 2-way ANOVA followed by Bonferroni’s multiple comparison test. Data are presented as scatter plus mean ± SEM. (n = 3 (WT 12mo), n = 4 (WT 12mo MR), n = 5 (rTg 12mo), n = 4 (rTg 12mo MR)). *p<0.05, **p<0.01, ***p<0.001, ****p<0.0001.

Next, levels of phosphorylated (pathological) tau in the brains of rTg4510 mice following MR intervention were examined in both soluble and insoluble protein fractions. In line with their transgenic make-up, rTg4510 animals displayed high protein levels of pTau at the PHF1 epitope for all bands compared to age-matched WT groups (genotype effect: F (1, 12) = 68.76, p<0.0001 for all, Fig. 3D1). No dietary effect was observed on PHF1 protein expression. However, protein analysis of total tau (Tau 5) demonstrated an overall diet effect (F (1, 12) = 10.52, p<0.01), with MR rTg4510 having higher levels compared with their control-fed littermates (Fig. 3D2). Similar changes were observed for the soluble CP13 epitope (Ser202), with increased tau phosphorylation in rTg4510 mice (genotype effect: F (1, 12) = 43.72, p<0.0001, Fig. 3E1). Confirming previous data, genotype (F (1, 12) = 90.73, p<0.0001) and diet (F (1, 12) = 6.720, p<0.05) effects for total tau brain expression was observed (Fig. 3E2).

The insoluble protein fraction was also investigated as it comprises highly aggregated tau. Corresponding to the soluble fractions, PHF1 insoluble protein expression was significantly higher in rTg4510 on both diets (all individual bands) (genotype effect: F (1, 10) = 23.72, p<0.001, Fig. 3G1). Dietary (F (1, 10) = 5.800, p<0.05) and genotype effects (F (1, 10) = 34.63, p<0.001) were observed for total tau insoluble protein levels from the PHF1 blots (Fig. 3G2). Next, the same analysis was conducted for the insoluble protein levels of CP13, with an overall genotype effect being observed (F (1, 12) = 36.88, p<0.001), following the same protein expression as the soluble fractions.

### Increased hippocampal inflammatory markers in rTg4510 mice on MR

To determine the effects of dietary MR on neuroinflammation, a hallmark of brain pathology in dementia, we examined the expression of glial inflammatory markers in the hippocampus. Astrocyte marker GFAP mRNA levels were increased in rTg4510 compared to WT mice (genotype effect: F (1, 26) = 159.5, p<0.0001), with no significant changes due to diet (Fig. 4A). Comparable results were obtained for CD68, with rTg4510 mice expressing increased levels of this microglial activation marker compared to WT (F (1, 26) = 51.08, p<0.0001, Fig. 4C). Additionally, Iba1 (microglia) and NLRP3 (inflammasome) gene expression revealed not only genotype effects (Iba1: F (1, 26) = 24.56, p<0.0001, NLRP3: F (1, 26) = 15.88, p<0.001, Fig. 4B & D) but also significant increases due to diet in rTg5410 mice (Iba1: F (1, 26) = 5.828, p<0.05, NLRP3: F (1, 26) = 8.990, p<0.01; Fig. 3B & D). An overall gene x diet interaction was also seen (Iba1: F (1, 26) = 5.285, p<0.05, NLRP3: F (1, 26) = 6.928, p<0.05), as no dietary alterations were detected in WT mice (Fig. 4B & D).

**FIGURE 4:**
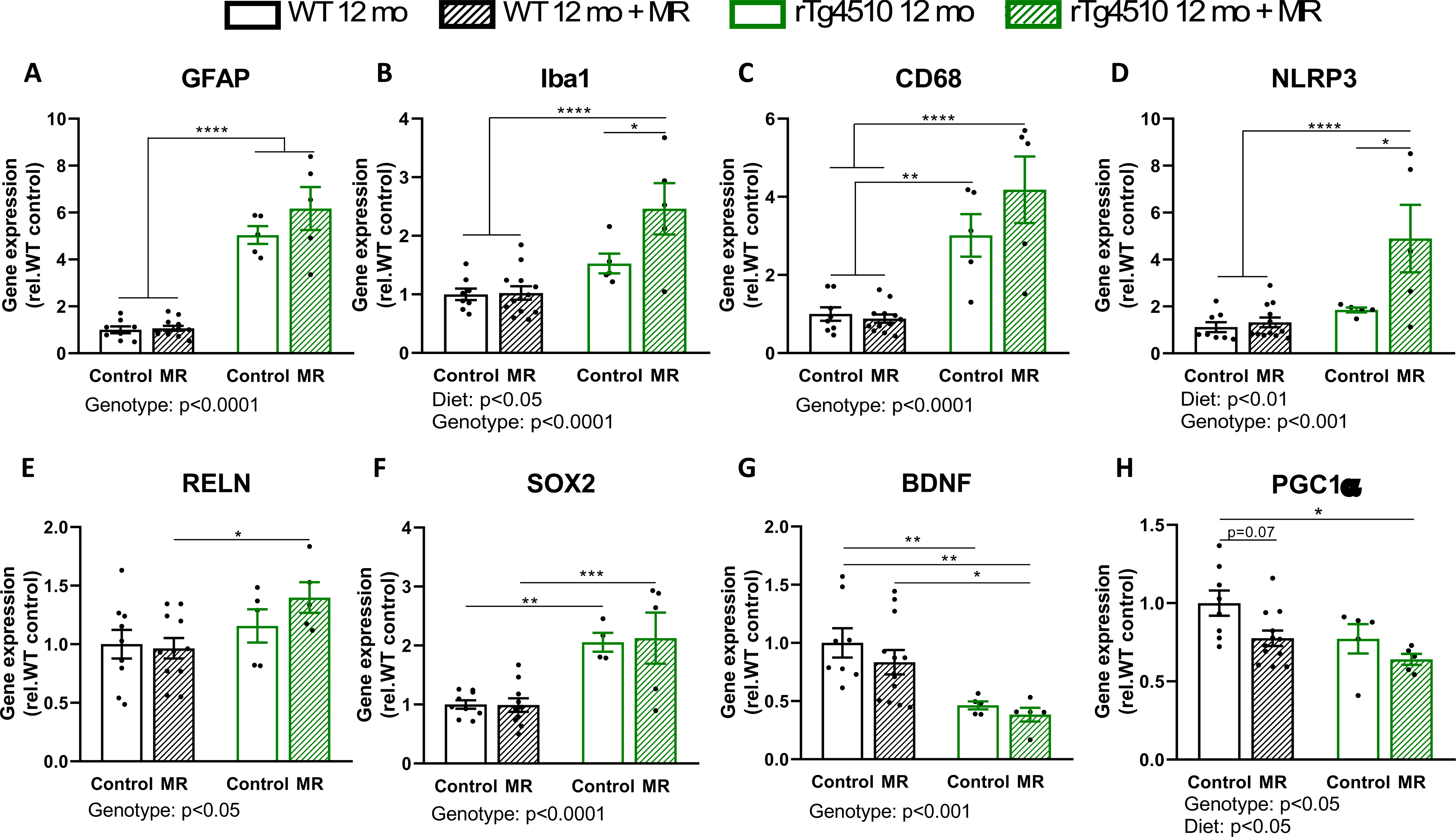
Neuroinflammation, neurogenesis, and neurotrophic markers in 12-month-old mice. Relative gene expression of GFAP, (B) Iba1, (C) CD68, (D) NLRP3, (E) RELN, (F) SOX2, (G) BDNF, and (H) Pgc1α in the hippocampus. YWHAZ was used as housekeeping gene. All data are shown as relative to the group WT (-/-) control diet. Data analysed by 2-way ANOVA followed by Bonferroni’s multiple comparison test. Data are presented as scatter plus mean ± SEM. (n = 8-9 (WT 12mo), n = 12 (WT 12mo MR), n = 5 (rTg 12mo), n = 5 (rTg 12mo MR)). *p<0.05, **p<0.01, ***p<0.001, ****p<0.0001.

Further investigating MR effects in the hippocampus, we observed elevated expression of plasticity and homeostasis mediators such as RELN, encoding for the extracellular matrix glycoprotein reelin (F (1, 26) = 5.641, p<0.05) and stem cell marker SOX2 (F (1, 24) = 30.97, p<0.0001, Fig. 4E & F), but low expression of nerve growth factor BDNF (F (1, 26) = 17.24, p<0.001) and mitochondrial regulator Pgc1α (F (1, 26) = 6.328, p<0.05) in rTg4510 vs. WT animals (overall genotype effects, Fig. 4G & H). These results suggest a tau-dependent modulation of neurogenesis and neuronal growth. While rTg4510 on MR displayed a significant increase in RELN and SOX2 expression compared to controls on the MR diet (Fig. 4E & F), an additional dietary effect (F (1, 26) = 6.106, p<0.05) was revealed for PCG1α expression (Fig. 4H).

Synaptic modifications were further analysed with the postsynaptic marker PSD95 and presynaptic protein synaptophysin. No significant differences were detected, revealing neither diet nor genotype effects (S4).

### Dietary MR increases glycolytic markers in the hippocampus of rTg4510 mice

Several mechanisms relevant for glucose homeostasis have been linked to neurodegeneration in tauopathies, such as impaired insulin signalling, mitochondrial function, and decreased glycolysis (24). We therefore investigated effects of diet and genotype on the glycolytic pathway in the hippocampus by determining gene expression of key regulatory markers of glycolysis (Fig. 5A). MR significantly increased hexokinase 2 (HK2) but not HK1 gene expression (HK2 diet effect: F (1, 27) = 4.848, p<0.05), alongside a genotype effect (F (1, 27) = 104.2, p<0.0001), where MR rTg4510 displayed higher levels compared to other groups (Fig. 5B, C). A diet x genotype interaction (F (1, 27) = 18.43, p<0.001) and overall genotype effect (F (1, 27) = 10.40, p<0.01) were also observed for glyceraldehyde 3-phosphate dehydrogenase (GAPDH), which was significantly lower in rTg4510 mice on the control diet compared to WT (Fig. 5D). MR rescued its expression in rTg4510 animals to control levels (Fig. 5D). Furthermore, the MR effect was different in WT mice, with decreased GAPDH expression compared to WT on the control diet (Fig. 5D). A similar effect was seen for phosphoglycerate dehydrogenase (Phdgh) expression; an overall interaction (F (1, 24) = 6.615, p<0.05) and genotype effect (F (1, 24) = 17.68, p<0.001) were observed (Fig. 5E). rTg4510 mice had higher expression of Phdgh compared to WT mice, and even higher levels were found in transgenic animals on MR. Here, no significant dietary effect was observed in WT mice (Fig. 5E). Following on, pyruvate kinase (PK) was significantly higher in rTg4510 mice compared to WT (genotype effect: F (1, 26) = 5.848, p<0.05), with normalisation vs WT achieved by MR (Fig. 5F).

**FIGURE 5:**
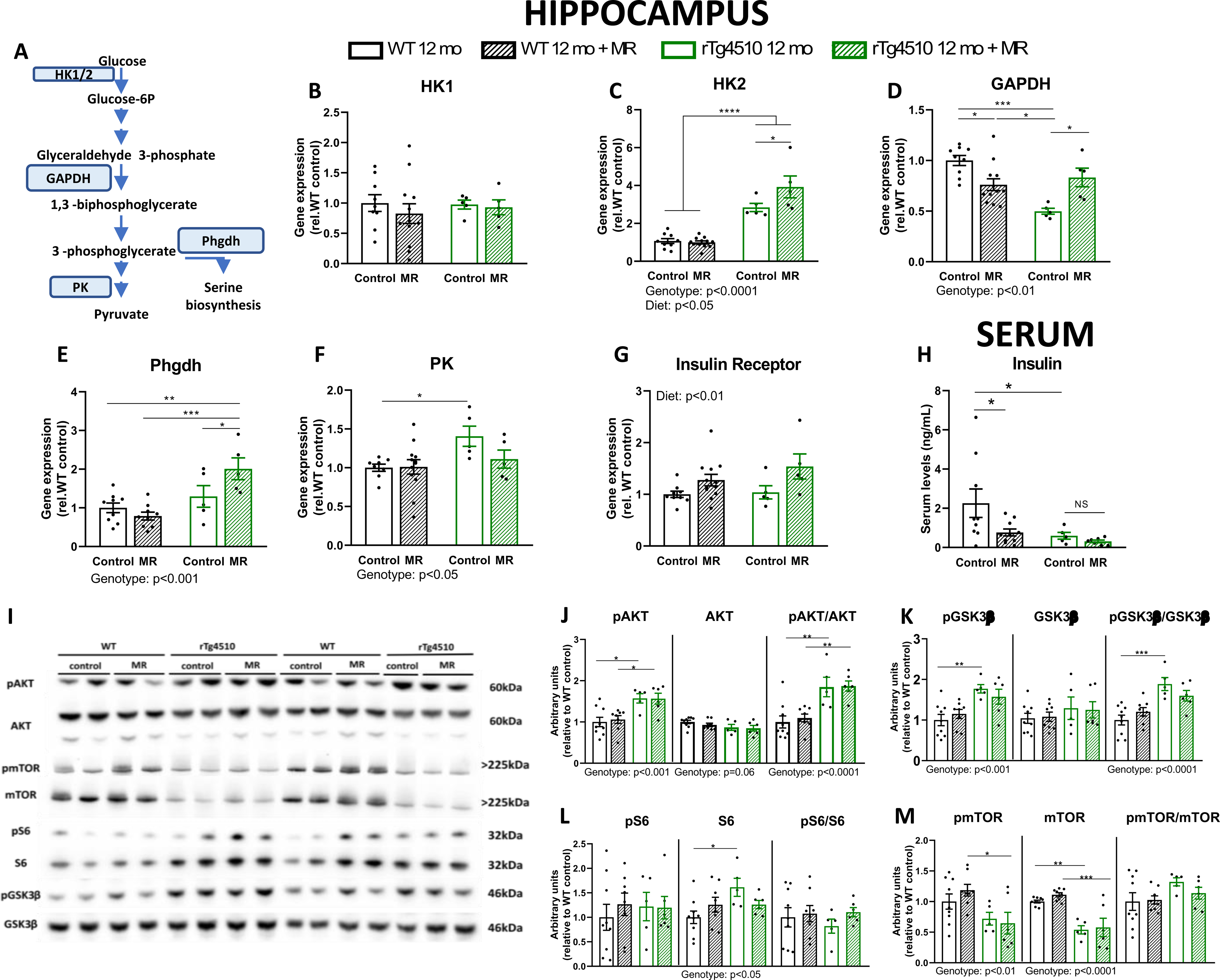
Glucose metabolism and insulin signalling in the brain of 12-month-old mice. (A) Short schematic representation of the glycolysis pathway. Relative gene expression of (B) HK1, (C) HK2, (D) GAPDH, (E) Phgdh, (F) PK, and (G) IR (n = 9 (WT 12mo), n = 12 (WT 12mo MR), n = 5 (rTg 12mo), n = 5 (rTg 12mo MR)). (H) Insulin serum levels (ng/mL) (n = 9 (WT 12mo), n = 10 (WT 12mo MR), n = 5 (rTg 12mo), n = 7 (rTg 12mo MR)). (I) Representative immunoblots of pAKT (Ser473), total AKT, pmTOR (Ser2448), total mTOR, pS6 (Ser235/237), total S6, pGSK3β (Ser9), and total GSK3β. (J) Densitometry analysis of pAKT, total AKT, and the ratio between phosphorylated and total AKT. (K) Densitometry analysis of pGSK3β, total GSK3β, and the ratio between phosphorylated and total GSK3β. (L) Densitometry analysis of pS6, total S6, and the ratio between phosphorylated and total S6. (M) Densitometry analysis of pmTOR, total mTOR, and the ratio between phosphorylated and total mTOR. YWHAZ was used as housekeeping gene and data plotted as relative to WT control diet group for qPCR data. All Western Blot data were normalised by Ponceau as loading control and plotted as relative to WT control diet group. Analysis of phosphorylated, total, and phospo/total ratio were individually performed and plotted in the same graph for visualisation purposes. (n = 9 (WT 12mo), n = 12 (WT 12mo MR), n = 5 (rTg 12mo), n = 6 (rTg 12mo MR)). Data analysed by 2-way ANOVA followed by Bonferroni’s multiple comparison test. Data are presented as scatter plus mean ± SEM. *p<0.05, **p<0.01, ***p<0.001, ****p<0.0001.

To determine systemic insulin signalling, we first determined fasted serum insulin levels. As expected, MR significantly decreased fasted serum insulin levels in WT animals (15,33) (Fig. 5H). rTg4510 mice on the control diet already had significantly lower levels of serum insulin compared to WT on the same diet, comparable to WT MR levels (Fig. 5H). No dietary effect was observed in rTg4510 mice (Fig. 5H). Next, the effects of tau transgene and diet on brain insulin signalling were determined. MR significantly increased total insulin receptor (IR) levels in the hippocampus of both, WT and rTg4510 (overall diet effect: F (1, 27) = 7.983, p<0.01, Fig. 5G). Next, protein levels of relevant proteins in the insulin signalling pathway were determined by Western Blotting (Fig. 5I), with phosphorylated AKT and GSK3β being significantly higher in rTg4510 compared to WT mice in both groups (overall genotype effect; pAKT: F (1, 24) = 20.97, p<0.001, pGSK3β: F (1, 24) = 17.16, p<0.001, Fig. 5J, K). No significant changes were observed for the total protein levels of AKT and GSK3β, resulting in increased relative phosphorylation levels of both proteins in rTg4510 mice (overall genotype effect; pAKT/total: F (1, 24) = 28.83, p<0.0001, pGSK3β/total: F (1, 24) = 24.47, p<0.0001, Fig. 5J, K). Lower levels of pmTOR (Ser2448) were detected in rTg4510 vs controls (genotype effect: F (1, 24) = 9.639, p<0.01), similar to total mTOR levels (genotype effect: F (1, 24) = 45.79, p<0.0001, Fig. 5M). No differences were detected for the pmTOR/total ratio (Fig. 5M). There were no significant changes in pS6 or pS6/total ratio (Fig. 5L). Nevertheless, rTg4510 mice had increased total S6 levels compared to WTs, with a trend towards a decrease in the MR group (rTg4510 control vs MR: p=0.09, Fig. 5L) as indicated by an overall genotype effect (F (1, 24) = 4.494, p<0.05) and interaction (F (1, 24) = 4.378, p<0.05; Fig. 5L).

### MR increases serum FGF21 and hippocampal FGFR1 expression

In line with MR’s reported action to increase levels of circulating FGF21 (15), an overall dietary effect (F (1, 26) = 22.93, p<0.0001) was observed for serum levels at 12 months (Fig. 6A). Notably, an age effect was also detected (F (1, 19) = 4.431, p<0.05), with older WT mice displaying significantly lower FGF21 serum levels. There were no differences between 6– and 12-months rTg4510 mice (Fig. 6A), yet FGF21 gene expression in the brain was significantly higher in rTg4510 compared to WT mice at 6 months. No differences were seen for the FGF21 co-receptor klotho (Fig. 6B, C).

**FIGURE 6:**
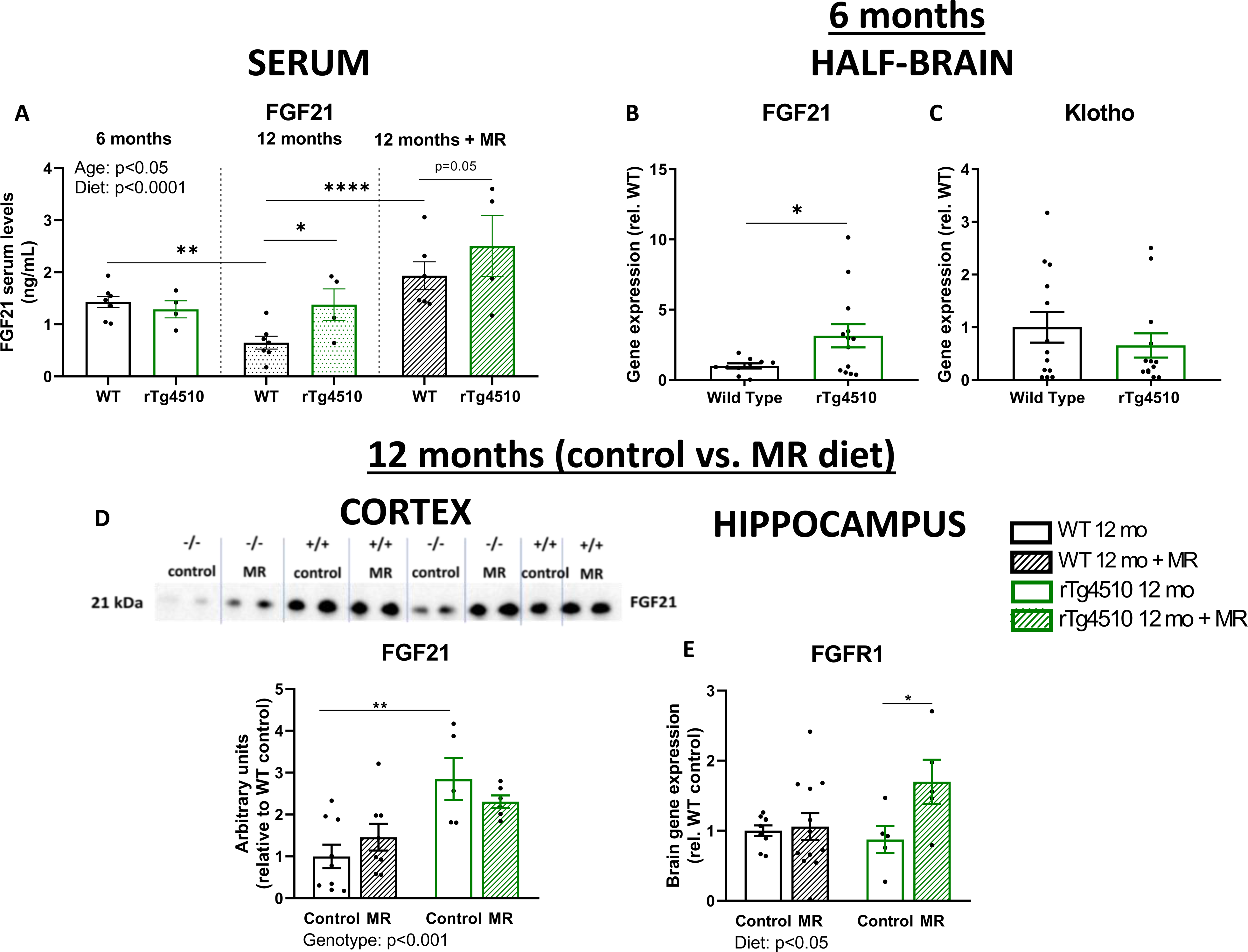
FGF21 signalling. (A) Serum levels of FGF21 (ng/mL) (n = 8 (WT 6mo), n = 4 (rTg 6mo), n = 11 (WT 12mo), n = 11 (WT 12mo MR), n = 4 (rTg 12mo), n = 4 (rTg 12mo MR)). Age (WT vs rTg 6 months) and diet (control vs MR 12 months) analyses were performed separately by 2-way ANOVA followed by Bonferroni’s multiple comparison test. Data were plotted together for visualisation purposes and presented as scatter plus mean ± SEM. (B, C) Relative FGF21 and Klotho gene expression in the brain of 6-month-old mice (n = 12-13 (WT 6mo), n = 12-13 (rTg 6mo)). (D) Representative immunoblots and densitometric analysis of FGF21 protein levels in the cortex of 12-month-old mice. Data were normalised by Ponceau as loading control and plotted as relative to WT control group. (n = 9 (WT 12mo), n = 8 (WT 12mo MR), n = 5 (rTg 12mo), n = 6 (rTg 12mo MR)). (E) Hippocampal FGFR1 gene expression. YWHAZ was used as housekeeping gene and data plotted as relative to WT control group. (n = 9 (WT 12mo), n = 9 (WT 12mo MR), n = 5 (rTg 12mo), n = 5 (rTg 12mo MR)). Data analysed by 2-way ANOVA followed by Bonferroni’s multiple comparison test. Data are presented as scatter plus mean ± SEM. *p<0.05, **p<0.01, ***p<0.001, ****p<0.0001.

We next evaluated protein levels of FGF21 in the cortex of WT and rTg4510 after MR (Fig. 6D). rTg4510 displayed higher levels compared to WT littermates (genotype effect: F (1, 24) = 16.87, p<0.001). No dietary effect was detected (Fig. 6D). Nevertheless, FGFR1 gene expression in the hippocampus of rTg4510 was significantly increased by MR (F (1, 27) = 4.408, p<0.05, Fig. 6E), mirroring FGF21 serum levels (Fig. 6A). This dietary effect was not observed in WT mice, suggesting a genotype-dependent effect of the diet (Fig. 6E).

## Discussion

This work aimed to determine the behavioural and cognitive phenotypes of rTg4510 in an age-dependent manner and in response to dietary MR. The diet improved short-term memory and habituation, as well as motor coordination and social recognition in rTg4510 mice. Increased metabolic markers associated with glucose utilisation and FGF21 signalling may play a role in the beneficial behavioural phenotype observed.

Our understanding of metabolic changes in FTD remain incomplete, especially in females, which display more pronounced tauopathy (27,28). In our study, rTg4510 mice had lower body weight and fat mass in comparison to control mice despite spending more time in water and food zones. Previous MR protocols uncovered changes in body composition in other animal models (34–38). Here, MR caused a specific reduction in fat mass only in WT mice, suggesting differing metabolism effects cf. rTg4510 mice (15,17).

A typical trait of the rTg4510 model is hyperactivity, detected as early as 2 months of age (39). This was maintained beyond habituation at 6 months of age in our study. The continuous hyperactivity of rTg4510 mice compared to WT aligns with disturbances in the sleep-wake cycle and clock gene observed in rTg4510 mice, alongside with a tau-dependent phenotype (40). Furthermore, rTg4510 mice on MR demonstrated exacerbated hyperactivity compared to control diet. MR, like CR, has been known to increase locomotor activity in mice (14,15,41). Higher energy expenditure suggests a correlation with hyperactivity, increased metabolic rate, and lower body weight (33,42). However, age, not hyperactivity, correlated with body weight in the rTg4510 model (43). In contrast to our findings, a basic CR diet did not impact hyperactivity in rTg4510 animals, indicating diverse dietary effects between CR and MR (42).

Deficits in motor coordination is another trait of rTg4510 mice observed in our study. Divergence from previous studies may be attributable to gender differences (29,44), but important to note that such deficits correlate with tauopathies in different FTD models, with females exhibiting a more severe pathology (45,46). Here, motor learning was only dramatically affected at 12 months of age, implying the progressive degeneration of brain areas involved. Our study also revealed an important MR effect on improving rTg4510 motor coordination and learning, which was not seen in the WT group. Similarly, previous research did not report differences in motor coordination in C57BL/6J male mice on MR (13). Interestingly, rTg4510 on CR did not exhibit improvements in motor coordination after 12 weeks of diet (42), but males and females were pooled, and CR started at 4 months of age (42). MR and CR therefore differ again regarding motor coordination in the rTg4510 model, though CR has previously been shown to decrease age-related impairment of motor performance in rodent models (47,48).

Previous CR studies suggested cognitive benefits in aged mice (49), similar to our results revealing MR related improvements in short-term spatial memory in both, rTg4510 and WT mice at 12 months. Further corroboration can be found in recent studies indicating that 3 months of MR improved age-related short– and long-term spatial memory in WT mice, demonstrated using Y-maze and water maze (WM) tasks (22). More recently, MR was shown to improve long-term spatial memory in middle-aged insulin-resistant mice by reducing oxidative stress and neuronal death in the hippocampus (50). Early CR studies emphasized the role of dietary intervention in improving spatial memory and hippocampal plasticity in rodents (49), more specific in increasing the number of dividing cells in the hippocampus of aged female mice, but not in males (51). However, rTg4510 mice on CR showed no improvement in WM (42), indicating the need for cautious interpretation based on transgenic mutation, sex, and dietary approach (e.g., CR vs MR)

The OF-NO-SI task affirmed MR’s positive impact on habituation and memory deficits in the rTg4510 model, especially in social interaction. This study represents the first evaluation of short-term and recognition memory in rTg4510 mice with MR intervention, measuring both activity and cognition within the same paradigm (52). Data exist on the evaluation of these tasks separately, spread over different days (14,22,42,53), also reporting reduced interest in social interaction (42), while adult male rats on 25% and 50% CR showed greater social exploration (54). Previously, rTg4510 mice exhibited significant deficits in long-but not short-term novel object recognition memory (55), in line with our study.

As previously reported (26), rTg4510 mice displayed robust hTau gene expression in the hippocampus, but no alterations in endogenous mouse tau. On MR, a small but significant increase in hTau gene expression was detected, alongside higher total soluble tau protein levels in rTg4510 mice. Notably, the rise in total non-phosphorylated tau, particularly in the soluble fraction, could enhance normal function and improve cognition by interacting with tubulin (56). Indeed, SantaCruz et al. (2005) discovered that suppressing tau gene transcription with doxycycline did not halt pTau accumulation in neocortex and hippocampus in this model. However, cognitive improvement was observed, aligning with our findings (26).

Neuroinflammation, marked by increased expression of brain GFAP (astrocyte marker) and Iba1 (microglial marker), was previously detected in rTg4510 as early as 2.5 months of age, highlighting another mechanism matching FTD pathology (57). rTg4510 presented with high expression of GFAP and CD68 levels in hippocampal tissue, without alterations by MR. This contrasts with prior research suggesting that MR improves cognitive impairments by decreasing inflammation in the brain (23). Expression of Iba1 and NLRP3 were significantly higher in the MR rTg4510, which aligns with the raised total Tau levels, suggesting increased microglial activation. While protein aggregation and microglial activation can exacerbate each other via proinflammatory cytokines such as TNFα and IL-1β (58), microglial activation is equally crucial for clearance and mitigation of tau toxicity (59). The interplay of these factors in connection with MR require further investigation ideally involving multiple time points to identify cause and effect.

Recent findings show adult neurogenesis persists but declines with age, impacted by neurodegeneration (60). Surprisingly, rTg4510 mice exhibit increased RELN and SOX2 gene expression in the hippocampus. Neurotrophic factors BDNF and PGC1α decrease, potentially triggering compensatory heightened neurogenesis (61). In contrast, chronic methionine treatment reduces neurogenesis in mice (62), but our model reveals no opposite effects on neurogenesis with MR.

Hyperphosphorylation of tau can affect glucose metabolism in the brain, with impairments contributing to tauopathies (63), which can directly inhibit aerobic glycolysis and increase insulin resistance (64,65). We observed low gene expression of GAPDH, a key glycolytic enzyme, in the brains of rTg4510, followed by restoration of its expression in FTD mice on MR. Our results are in line with low cerebral glucose metabolism observed in FTD (66). It is proposed that pathological tau strongly interact with GAPDH, affecting glycolysis and apoptosis (67), and that low GAPDH levels may act protectively by redirecting glucose metabolism to the pentose phosphate pathway (68). In addition, increased HK2 and Phgdh in FTD mice on MR may therefore activate glycolysis in these mice, promoting serine metabolism and potentially contributing to the observed cognitive improvements. The role of glycolytic enzymes in spatial memory is important for memory formation as previously described (69), but further investigation is needed.

The mechanism of MR improving hippocampal-dependent memory is not fully understood, but significantly improved spatial cognition in aged mice was associated with increased hepatic and serum FGF21 levels, with FGF21 knockdown lowering MR benefits on cognition (23). The neuroprotective role of FGF21 was linked with reduced apoptosis and tau hyperphosphorylation (70). In our study, ageing lowered FGF21 serum in WT mice only and dietary MR successfully increased levels in both genotypes. Notably, raised brain FGF21 expression was found in 6-month rTg4510 mice, levels were also higher in serum and cortex of rTg4510 mice at 12 months, while expression of FGFR1 increased in the rTg4510 MR group.

In summary, our results demonstrate that neuronal overexpression of mutated human Tau in mice causes behavioural impairments alongside inflammation and alterations in brain glucose metabolism. After an 8-week dietary MR, 12-month-old female rTg4510 mice exhibited cognitive and behavioural improvements, associated with increased aerobic glycolytic metabolism and FGF21 signalling. These effects were independent of cortical pTau expression or alterations in neurogenesis or whole-body composition.

## Methods

### Animals

The study has been approved by the University of Aberdeen Ethics Review Committee Board and the UK Home Office in accordance with the Animal Scientific Procedures Act 1986 and ARRIVE 2.0 guidelines (71). rTg4510 animals were purchased from The Jackson Laboratory (RRID: IMSR_JAX:024854, FVBxC57BL/6J background) and bred at Charles River Laboratories, UK, Mice were transferred to the Medical Research Facility (MRF) at the University at 6 months of age with at least 2 weeks of habituation prior to experimentation. The rTg4510 mice were created by crossing mice carrying the human MAPT^P301L^ cDNA downstream of a tetracycline operon-responsive element (tetO) to animals expressing a tetracycline-controlled transactivator (tTA), which is controlled by a calcium/calmodulin-dependent kinase II (CaMKIIa) promoter (26). The human tau^P301L^ transgene is constitutively expressed in bi-transgenic offspring, which can be inactivated by administration of the tetracycline analogue doxycycline. The MAPT^P301L^ transgene encodes human tau with the P301L mutation. All rTg4510 mice used in this study are bi-transgenic for tetO and tTA (+/+), while WT littermates are negative for both genes (-/-). Genotype confirmation was performed by Transnetyx Inc. using ear-biopsies.

6-month-old female mice were group housed (individually housed if specified in the protocol) and exposed to a 12h light/dark cycle (7am – 7pm) in a climate-controlled environment (20 – 22°C, 60 – 65% humidity) with ad libitum access to food and water. Enrichment was provided with paper strips and cardboard tubes. Animals were grouped by genotype and/or body weight, and randomisation was performed using a random number generator (www.random.org) with the experimental IDs before in vivo tests.

In vivo tests were conducted at 6 months of age and animals aged until 10 months, when groups were split into control (0.86% DL-methionine) and MR diet (0.172% DL-methionine) (Dyets, USA) for 8 weeks. The last 4 weeks of MR intervention were accompanied by in vivo tests, with mice being 12 months-old at the end of the study. The experimental timeline and total number of mice use per replication are detailed in S1. Study termination involved a 5-hour fast, humane culling by cervical dislocation. Trunk blood was collected for serum into separator micro-tubes (BD Microtainer), brains snap-frozen in liquid nitrogen, stored at –80°C. Blinding wasn’t possible due to rTg4510 mice hyperactivity and diet colours.

### PhenoTyper

Locomotor and circadian activity were assessed using the PhenoTyper home cages (Noldus) as previously described (72). PhenoTypers are clear Perspex boxes (30 x 30 x 45 cm W x L x H) with a lid carrying the infrared system and infrared sensitive camera for continuous recording. Singly housed animal was continuously tracked (25Hz sampling rate) at the centre of gravity by background subtraction for a total of 168 hours during seven consecutive days by Ethovision 11.5 (Noldus). Analysis was carried out using Ethovision software. The first 3 hours were used to determine habituation to a new environment. Circadian and motor activity was evaluated using the hourly average of distance moved over 5 days (Sunday at noon – Friday at noon). Average activity over 24-hour periods was pooled per hour, and time in food and water zone determined for both light and dark periods. Food, water, and mice were weighed before and after each test. A maximum of 24 cages were recorded simultaneously, each mouse single-housed throughout. Cages were washed after each experiment.

### RotaRod

Motor coordination and motor learning were evaluated using the RotaRod apparatus for mice (UgoBasile NG RotaRod) (73). In brief, the apparatus consists of 4 parallel rotating rods, separated by a wall. Two identical RotaRods were used side by side in a dedicated experimental laboratory with overhead ambient lighting (100 lux) and a temperature of ±21°C. The tests were performed on 2 consecutive days with 4 trials/day, on acceleration mode from 5 – 45 rpm up to 300 seconds. Animals were randomised using a Latin square design and taken to the experimentation room 30 minutes before the test for habituation. They were placed on the rod during a low rotation (5 rpm) and acceleration was started once all 4 mice were placed on their rods and progressed with a forward motion. Trials terminated when the mice fell off the rod or at the maximum time of 5 minutes; 2 min-between-trial intervals were given. Rods were cleaned between trials with fragrance-free wipes. Motor learning was determined by the total time spent on the rod during trial 1 compared to trial 8, and motor coordination assessed by latency to fall per trial.

### Y-maze

Spatial working memory and spontaneous alternations were determined using the Y-maze test (74). The Y-maze apparatus (Fig. 2A) is made of Perspex and consists of 3 arms protected by walls (60cm L, 10cm H), labelled A, B, and C with indirect lights (175 lux). Animals were, habituated to the novel environment for 30 minutes and then placed into the distal end of the start arm B facing the wall. Subjects were allowed to explore the maze for 10 minutes and their activity was tracked as centre of gravity subtracting their background by an overhead camera at a sampling rate of 25 Hz connected to the Ethovision 11.5 software (Noldus). Triplet sequences (e.g., ABC, BCA, CAB) were used to determine the percentage of correct and incorrect alternations. Correct alternations comprised a triplet sequence visiting all three arms. Repeating an arm in a triplet sequence deemed alternation incorrect.

### OF-NO-SI paradigm

The Open Field-Novel Object-Social Interaction (OF-NO-SI) paradigm (31,52) assessed behaviour in novel (trial 1) and familiar environment (trial 2). Animals, acclimated for 30 minutes in controlled conditions (75 lux, ±21°C room temperature), were tracked in a 50cm diameter white Perspex circular arena with an overhead camera (Sony) (background subtraction, sampling rate 15Hz). ANY-Maze video tracking software (Ugo Basile SRL) analysed ambulatory activity. The tests were performed as described (31,52), divided into open field (OF), novel object (NO), and social interaction (SI), each with two 10-minute trials and a 15-minute inter-trial interval (ITI). Animals were individually placed into the arena for the first OF trial (OF1, novel environment) and continuously recorded. Next, mice were placed into their transfer cages (ITI) before the second OF trial (OF2) commenced (familiar environment). Once completed, a small wire cage cylinder (object: 8 cm diameter, 20 cm tall) was placed into the centre of the arena, and NO trials 1 and 2 were performed as described for OF. Subsequently, a sex and age-matched stranger mouse was introduced into the wire cage cylinder for interaction trials SI1 and SI2 as per previous segments. Stranger mice were housed in a different room than the experimental mice, and only brought to the room for testing. Stranger mice were rotated daily to minimize stress, and a maximum of 4 experiments per stranger mouse was performed on a same day. Animals were placed close to the wall of the arena at the start of each trial and the arena was cleaned with fragrance-free wipes at the end of each trial. Total distance moved, time spent in the wall zone (<5cm from wall of the arena), and time spent in the interaction zones (NO or SI, <5cm from cylinder) were obtained from the ANY-maze software for analysis (6.0, Ugo Basile).

### Echo MRI

Whole body mass composition was measured using Magnetic Resonance Imaging (MRI) technology (EchoMRI) (73). Fat and lean mass were determined by placing non-anesthetized mouse into a tube in a prone position allowing only small movements and accurate scanning. Echo MRI was only performed in the replication 1 of this study.

### Glucose Tolerance Test (GTT)

Animals were fasted 5 hours prior to the test. Before the GTT, mice were randomised and placed in individual test cages. Tail fasting blood glucose (time 0) was obtained by tail puncture with a needle and measured using the AlphaTRAK II glucometer for small animals (Berkshire), followed by an intraperitoneal (i.p.) injection of glucose solution (20% w/v glucose) in the dose of 2mg/g body weight (b.w.). Tail blood glucose was measured at the times 15-, 30-, 60-, and 90-minutes post-injection, as described (75).

### RT-qPCR

RT-qPCR was conducted following established procedures (76). Briefly, RNA was extracted from frozen tissue using TRI Reagent (Trizol; Sigma-Aldrich). RNA concentration was determined by micro-volume spectrophotometry (Genova Plus), and cDNA synthesis was performed using a Tetro cDNA synthesis kit (Bioline) following the manufacturer’s instructions using the GS-1 G-Storm thermocycler (LabTech). Diluted cDNA samples (1:10 ratio) were used for gene expression analysis. Amplification of genes utilized GoTaq qPCR Master Mix (Promega) as per the manufacturer’s protocol on the LightCycler® II 480 system machine (Roche). Primer sequences are detailed in Table S2. Quantitative cycle (Cq) was computed, and gene expression was determined using the Pfaffl equation, normalized to the housekeeping gene Ywhaz. Housekeeping stability was assessed pre-target gene amplification using RefFinder (77).

### Immunoblotting

Immunoblotting was done as previously described (78,79). In brief, brain tissues were homogenised in NP40 lysis buffer (1M HEPES, 5M NaCl, 0.1 EDTA, 1% NP-40 (Sigma): pH = 7.6) with protease and PhosStop inhibitors (Roche). Soluble proteins were obtained after centrifugation (12000 x rpm, 4°C, 20 minutes). Insoluble Tau was detected by extracting fractions using 70% formic acid, followed by neutralization (4 parts of a neutralising buffer (2M Tris, 2M NaH2P04 in dH20). Protein quantification was performed using the BCA kit (Sigma), except for insoluble fractions due to due to formic acid’s denaturation effect. Instead, an equal volume of neutralised samples was loaded into the gels, and total protein loading was used to determine protein expression levels. Samples were separated by SDS-PAGE gel electrophoresis cell (Biorad) and proteins were transferred nitrocellulose membranes using transfer buffer (25 mM Tris, 192 mM glycine, 20% (v/v) methanol), confirmed by Ponceau-S staining, used also as protein-loaded control. Immunoblotting utilized primary and secondary antibodies (Table S2), and images were captured at 16-bit by a Fusion CCD camera (Peqlab) and analyzed using ImageJ software (NIH). Phosphorylated proteins were normalized to their total values. Ponceau-S-stained total loading control was used for quantification. Adjusted values were expressed relative to the control group for each experiment. Proteins normalised by the total protein-loaded control.

### Insulin and FGF21 ELISA

Enzyme-linked immunosorbent assays (ELISAs) were used to determine the levels of insulin and FGF21 in the serum (15,80), following the respective supplier’s instructions (insulin: Merck Millipore #EZRMI-13K and FGF21: R&D Systems #MF2100). The readings were measured by spectrophotometry using a FLUOstar® Omega multi-mode plate reader (BMG Labtech).

### Statistical Analysis

Statistical analysis was conducted using GraphPad Prism 8 software. 2-way ANOVA or 3-way ANOVA were performed for data sets with multiple factors followed by Bonferroni’s or Tukey’s multiple comparison post-tests, respectively. Analysis of habituation data for the PhenoTyper was performed by nonlinear regression (one-phase decay), followed by 3-way ANOVA (source of variations: time, genotype, age and/or diet). Effects of age and diet analyses were conducted separately but are represented in the same graph for visualisation purposes. Age analysis was performed between 6– and 12-month-old groups (WT and rTg4510, both on a control diet). Diet analysis was conducted between control and MR diet groups (WT and rTg4510, both 12 months old). All data sets were submitted to Grubbs’ outlier test before the final statistical analysis. Shapiro-Wilk test was also performed for assessment of normal distribution. Alpha was set to 0.05 and significance was assumed as p<0.05. Data are expressed as scatter plus mean ± SEM. Detailed statistical analyses for each set of experiments are described in the respective figure legends and in Supplementary Table 1.

## Acknowledgements

We acknowledge the financial support from MD current funding British Heart Foundation (PG/21/10555, FS/PHD/21/29105) and the Development Trust (AP2069). MSM received the Elphinstone Scholarship and the IMS studentship, both from the University of Aberdeen. We recognise the support with animal care and in vivo experiments provided by the staff of the MRF.

## Author contributions

MSM, BP, and MD designed the experiments. MSM performed the experiments with assistance from AS. MSM, AS, GR, BP, and MD contributed to the analysis and interpretation of data. MSM wrote the manuscript and AS, GR, BP, and MD critically revised the manuscript. The final version was approved by all authors.

