## Supplementary Figures for "Effects of age and dietary methionine restriction on cognitive and behavioural phenotypes in the rTg4510 model of frontotemporal dementia"

A

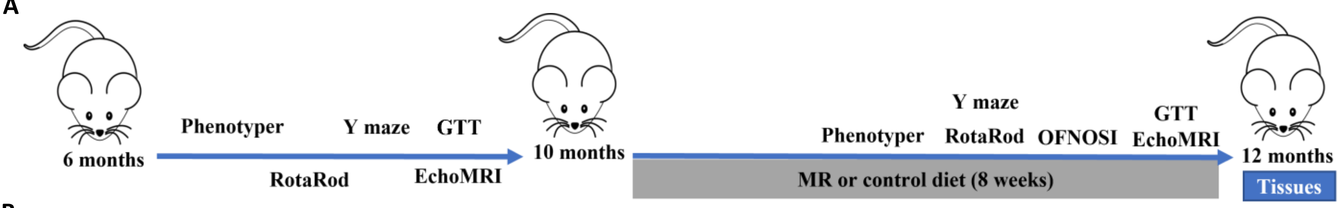


B

| Replication | Age | Dietary intervention | Wild Type (n) | rTg4510 (n) | Mice excluded | Age/group/reason of exclusion |
|---|---|---|---|---|---|---|
| **1** | 6 and 12 months | Control and MR | 9 | 11 | 1 | 10 mo/rTg + MR/weight loss |
| **2** | 6 and 12 months | Control and MR | 17 | 3 | 1 | 11 mo/rTg +MR/absent seizures, weight loss |
| **3 (tissues/blood collection)** | 6 months | control | 13 | 13 | - | - |

**Supplementary 1: Study design.** (A) Experimental timeline. 6-month-old mice performed a behavioural, cognitive, and metabolic test battery in the following order: PhenoTyper, RotaRod, Y maze, Echo MRI, GTT, and ITT. These mice aged until 10 months when half of the animals switched to an MR diet and the other half to a control diet for 8 weeks. In the last 4 weeks of MR intervention, PhenoTyper, RotaRod, Y maze, OFNOSI, Echo MRI, and GTT were conducted. At the end of the study, mice were 12 months old and humanly culled for tissues collection. (B) Number of mice used per replication**.** Total number of mice used at the start of each study at 6 months of age. 2 mice were humanly culled before the end of the study because of the healthy issues described in the table, and tissues from these mice were not used for molecular analyses. Endpoint of 20% body weight loss compared to the body weight at start of the study was used for exclusion.

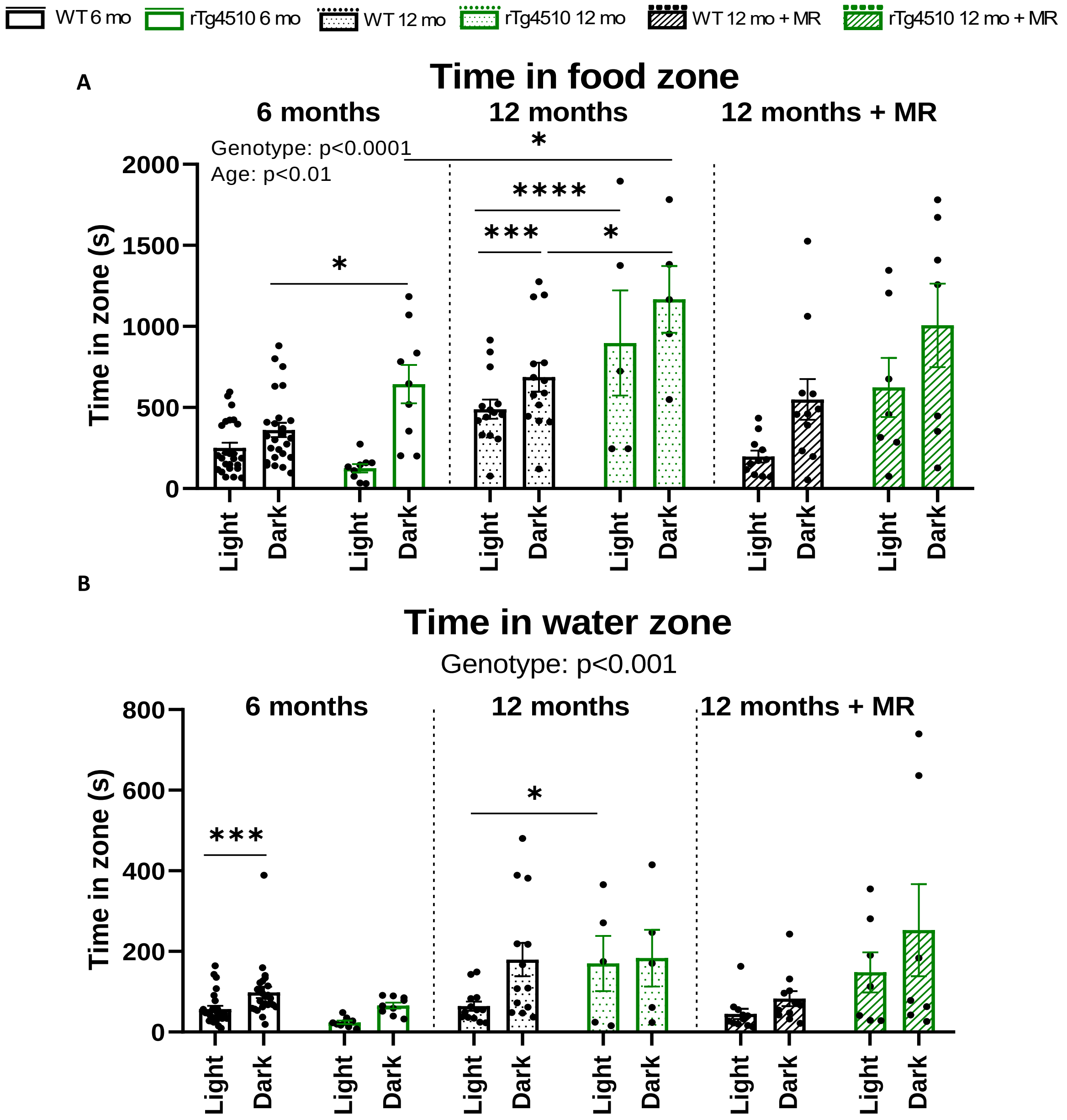


**Supplementary 2: Time in food and water zone during the PhenoTyper test.** (A) Time spent in food and (B) water zone during light and dark periods (n = 25 (WT 6mo), n = 12 – 14 (rTg 6mo), n = 9 (WT 12mo), n = 10 (WT 12mo MR), n = 5 (rTg 12mo), n = 7 (rTg 12mo MR)).

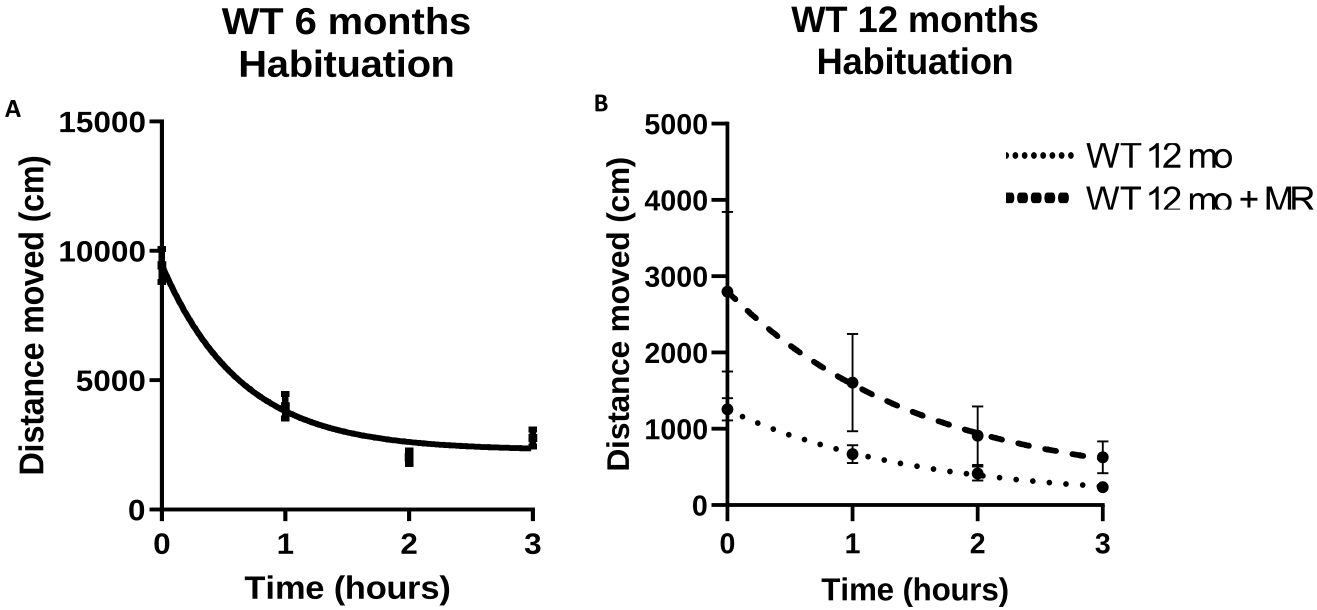


**Supplementary 3: WT habituation to a novel environment PhenoTyper home cage.** Distance moved (cm/hour) during 3h-habituation period at (A) 6 months of age, (B) 12 months of age in control and MR diet. Data are presented as scatter plus mean ± SEM. n = 25 (6 mo), n = 9 (control), n = 10 (MR). Graphs for visualisation purposes only.

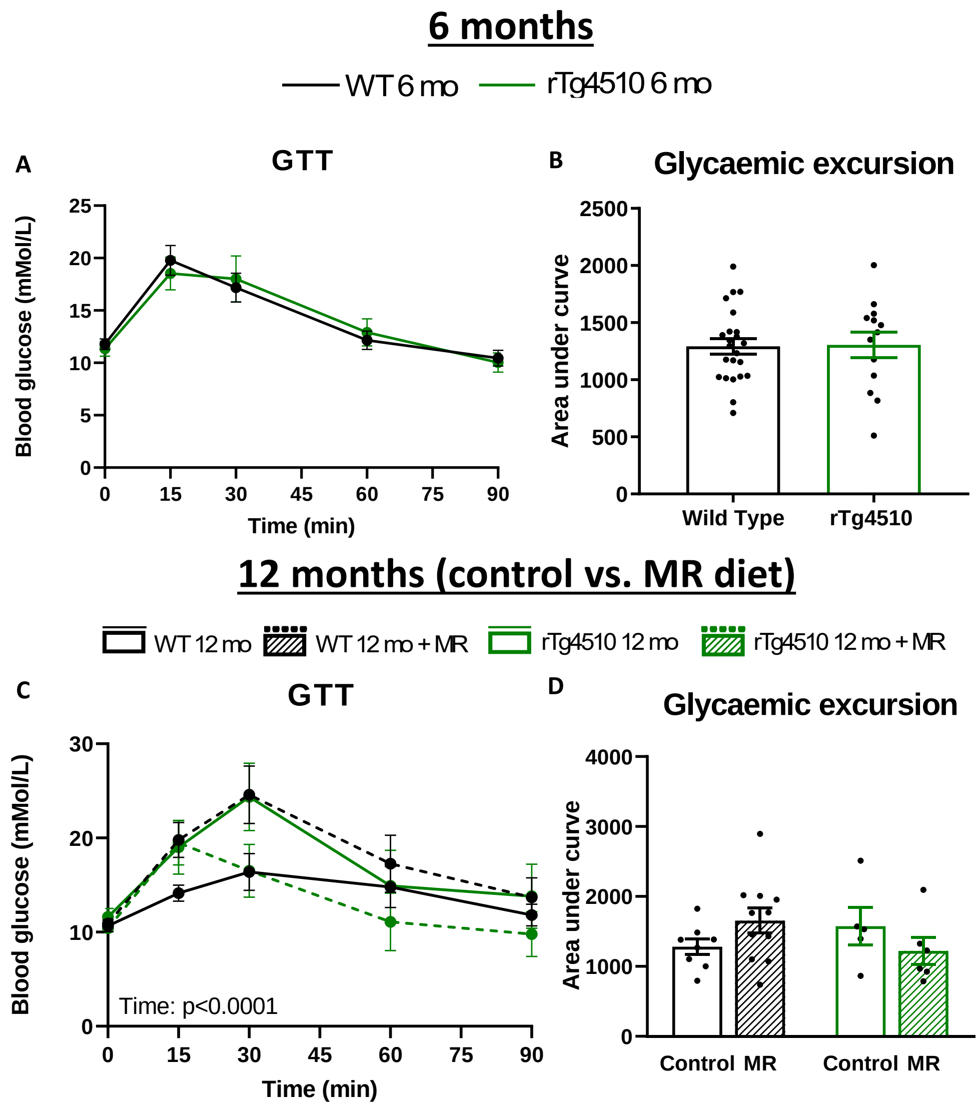


**Supplementary 4: Glucose tolerance test.** (A, C) Blood glucose levels vs. time during glucose tolerance test (GTT) and (B, D) area under the curve (AUC) analysis in 6- and 12-month-old animals. Data analysed by unpaired two-tailed t-test, 3-way ANOVA followed by Tukey's multiple comparison test or by 2-way ANOVA followed by Bonferroni's multiple comparison test. Data are presented as scatter plus mean ± SEM. n = 23 (WT 6mo), n = 13 (rTg 6 mo), n = 8 (WT 12 mo), n = 11 (WT 12 mo MR), n = 5 (rTg 12 mo), n = 6 (rTg 12 mo MR).

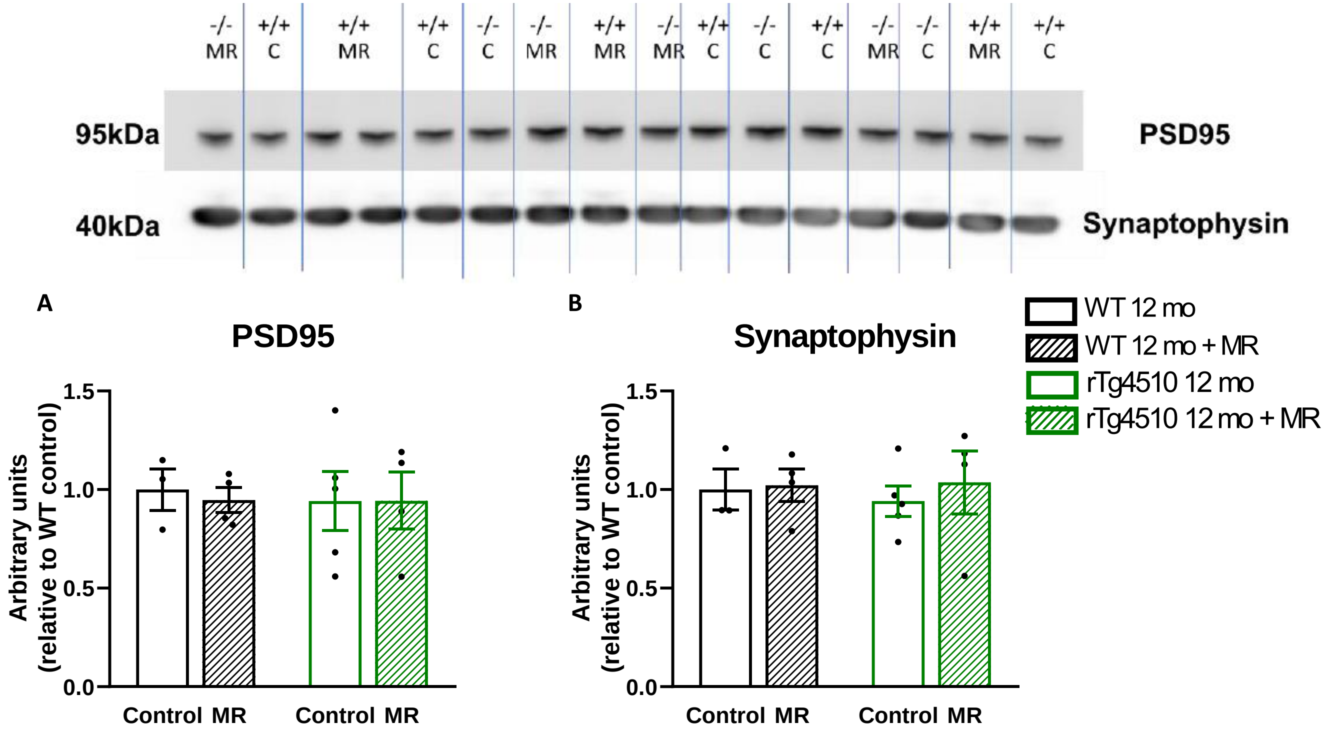


**Supplementary 4: Synaptic markers protein levels.** Representative immunoblots and densitometry analysis of (A) PSD95 and (B) synaptophysin in the cortex of 12-month-old mice. Data normalised by Ponceau as loading control and plotted as relative to the group WT (-/-) control diet. (n = 3 (WT 12mo), n = 4 (WT 12mo MR), n = 5 (rTg 12mo), n = 4 (rTg 12mo MR)). Data analysed by 2-way ANOVA followed by Bonferroni's multiple comparison test. Data are presented as scatter plus mean ± SEM.
