## Supplementary Tables for "Effects of age and dietary methionine restriction on cognitive and behavioural phenotypes in the rTg4510 model of frontotemporal dementia"

**Supplementary Table 1: List of statistically significant interactions**

|  | **Statistics** | | |
| --- | --- | --- | --- |
| **Figure** | **Analysis** | **Interaction (p value)** | **F (DFn, DFd)** |
| 1C | 2-way ANOVA | Interaction (p<0.05) | F (1, 37) = 5.165 |
| 1D | 3-way ANOVA | Genotype x Diet (p<0.05) | F (1, 111) = 6.157 |
| 1E |  | Phase effect (p<0.05) | F (1, 54) = 7.115 |
|  |  | Phase x Genotype (p<0.05) | F (1, 54) = 4.706 |
|  |  | Genotype x Diet (p<0.05) | F (1, 54) = 4.383 |
| 1F |  | Genotype x Diet (p<0.05) | F (1, 208) = 4.186 |
| 1G |  | Trial x Genotype (p<0.05) | F (1, 50) = 4.627 |
| 2B |  | Alternations x Genotype (p<0.01) | F (1, 54) = 8.045 |
|  |  | Alternations x Diet (p<0.0001) | F (1, 54) = 43.21 |
|  |  | Alternations x Genotype x Diet | F (1, 54) = 20.54 |
| 2D | Mixed-effects analysis | Trial x Diet (p<0.05) | F (1, 27) = 4.653 |
|  |  | Trial x Genotype x Diet (p<0.05) | F (1, 27) = 6.743 |
| 2F | 3-way ANOVA | Trial (p<0.0001) | F (1, 28) = 49.86 |
|  |  | Trial x Genotype | F (1, 28) = 7.063 |
| 2G | Mixed-effects analysis | Trial (p<0.0001) | F (1, 27) = 26.72 |
|  |  | Trial x Diet (p<0.01) | F (1, 27) = 3.122 |
| 3D1 | 2-way ANOVA | Interaction (p<0.05) | F (1, 12) = 6.698 |
| 3E2 |  |  | F (1, 12) = 5.156 |
| 4B |  |  | F (1, 26) = 5.285 |
| 4D |  |  | F (1, 26) = 6.928 |
| 5D |  | Interaction (p<0.001) | F (1, 27) = 18.43 |
| 5E |  | Interaction (p<0.05) | F (1, 24) = 6.615 |
| 5L (total S6) |  |  | F (1, 24) = 4.378 |
| Supplementary 2A | 3-way ANOVA | Phase effect (p<0.0001) | F (1, 49) = 38.54 |
|  |  | Phase x Age (p<0.01) | F (1, 49) = 7.366 |
|  |  | Genotype x Age (p<0.05) | F (1, 49) = 5.087 |
| Supplementary 4C |  | Genotype x Diet (p=0.0596) | F (1, 26) = 3.881 |

**Supplementary Table 2: List of primers used for qPCR**

| **Primers** | **Forward** | **Reverse** |
| --- | --- | --- |
| *human TAU* | CAGGAGTTCGAAGTGATGGAAGA | AGCCCCCCTGATCTTTCCT |
| *mouse Tau* | CAGGAGTTCGAAGTGATGGAAGA | AGCCCCCCTGATCTTTCCT |
| *GFAP* | CGGAGACGCATCACCTCTG | AGGGAGTGGAGGAGTCATTCG |
| *Iba1* | ATCAACAAGCAATTCCTCGATGA | CAGCATTCGCTTCAAGGACATA |
| *CD68* | CCTCGCCTAGTCCAAGGTC | GGATTCGGATTTGAATTTGGGCT |
| *NLRP3* | TGCTCTTCACTGCTATCAAGCCCT | ACAAGCCTTTGCTCCAGACCCTAT |
| *RELN* | TGAATGCAGCAACTGTGAGAT | ATCCAATCAGCGGTATTGTTCTT |
| *SOX2* | GCGGAGTGGAAACTTTTGTCC | GGGAAGCGTGTACTTATCCTTCT |
| *BDNF* | AGGTCTGACGACGACATCACT | CTTCGTTGGGCCGAACCTT |
| *PGC1α* | TATGGAGTGACATAGAGTGTGCT | GTCGCTACACCACTTCAATCC |
| *HK1* | CAAGAAATTACCCGTGGGATTCA | CAATGTTAGCGTCATAGTCCCC |
| *HK2* | ATGATCGCCTGCTTATTCACG | CGCCTAGAAATCTCCAGAAGGG |
| *GAPDH* | TGACCACAGTCCATGCCATC | GACGGACACATTGGGGGTAG |
| *Phgdh* | CCTCATTGTCCGGTCTGCTAC | CATCTTTCATCGAAGCTGTTGC |
| *PK* | GTGGCTCGGCTGAATTTCTCT | CACCGCAACAGGACGGTAG |
| *IR* | CCTGGTTATCTTCGAGATGGTCC | CCCCACATTCCTCGTTGTCA |
| *FGF21* | ACCTGGAGATCAGGGAGGAT | CACCCAGGATTTGAATGACC |
| *KLOTHO* | TCGGTACGTCTACTCACACCT | AGCCAACAAGTCTTTTTCCAGA |
| *FGFR1* | ACTCTGCGCTGGTTGAAAAAT | GGTGGCATAGCGAACCTTGTA |
| *YWHAZ* | GAAAAGTTCTTGATCCCCAATGC | TGTGACTGGTCCACAATTCCTT |

**Supplementary Table 3: List of antibodies and dilutions used for Western Blot**

|  | **Antibody** | **Dilution** | **Supplier** |
| --- | --- | --- | --- |
| **Primary antibodies** | PHF-1 (Tau p-Ser396/  p-Ser404) | 1:500 | P. Davies lab |
|  | CP13 (Tau p-Ser202) | 1:500 | P. Davies lab |
|  | AT5 (total tau) | 1:1000 | Abcam (#ab80579) |
|  | FGF21 | 1:1000 | Abcam (#ab171941) |
|  | p-AKT (Ser473) | 1:1000 | Cell Signalling Technology (#4060) |
|  | total AKT | 1:1000 | Cell Signalling Technology (#4691) |
|  | p-GSK3β (Ser9) | 1:1000 | Cell Signalling Technology (#9323) |
|  | total GSK3β | 1:1000 | Cell Signalling Technology (#9315) |
|  | p-S6 (Ser235/237) | 1:1000 | Cell Signalling Technology (#4858) |
|  | total S6 | 1:1000 | Cell Signalling Technology (#2217) |
|  | p-mTOR (Ser2448) | 1:1000 | Cell Signalling Technology (#2971) |
|  | total mTOR | 1:1000 | Cell Signalling Technology (#2983S) |
| **Secondary antibodies**  **(HRP conjugated)** | Mouse anti-rabbit | 1:10000 | Merck Millipore  (#AP188P) |
|  | Goat anti-mouse | 1:5000 | Merck Millipore (#AP181P) |
